# Transcriptomic signature and pro-osteoclastic secreted factors of abnormal bone marrow stromal cells in fibrous dysplasia

**DOI:** 10.1101/2024.02.23.581225

**Authors:** Zachary Michel, Layne N. Raborn, Tiahna Spencer, Kristen Pan, Daniel Martin, Kelly L. Roszko, Yan Wang, Pamela G. Robey, Michael T. Collins, Alison M. Boyce, Luis Fernandez de Castro Diaz

## Abstract

Fibrous dysplasia (FD) is a mosaic skeletal disorder caused by somatic activating variants in *GNAS*, encoding for Gα_s_, which leads to excessive cAMP signaling in bone marrow stromal cells (BMSCs). Despite advancements in our understanding of FD pathophysiology, the effect of Gα_s_ activation in the BMSC transcriptome remains unclear, as well as how this translates into their local influence in the lesional microenvironment. In this study, we analyzed changes induced by Gα_s_ activation in BMSC transcriptome and performed a comprehensive analysis of their production of cytokines and other secreted factors. We performed RNAseq of cultured BMSCs from patients with FD and healthy volunteers, and from an inducible mouse model of FD, and combined their transcriptomic profiles to build a robust FD BMSC genetic signature. Pathways related to Gα_s_ activation, cytokine signaling, and extracellular matrix deposition were identified. In addition, a comprehensive profile of their secreted cytokines and other factors was performed to identify modulation of several key factors we hypothesized to be involved in FD pathogenesis. We also screened circulating cytokines in a collection of plasma samples from patients with FD, finding positive correlations of several cytokines to their disease burden score, as well as to one another and bone turnover markers. Overall, these data support a pro-inflammatory, pro-osteoclastic behavior of BMSCs bearing hyperactive Gα_s_ variants, and point to several cytokines and other secreted factors as possible therapeutic targets and/or circulating biomarkers for FD.

## Introduction

Fibrous dysplasia (FD) is a mosaic skeletal disorder caused by somatic activating variants in *GNAS*, encoding the α subunit of the stimulatory G protein. The location and extent of bone lesions is variable and can be associated with hyperpigmented skin macules and/or various endocrinopathies, termed McCune-Albright syndrome (MAS, OMIM #174800). Gα_s_ p.R201C and p.R201H activating substitutions are typically the genetic cause, and a minority of patients carry a p.G227L variant. These mutations lock Gα_s_ in an active conformation, leading to excessive production of intracellular cAMP in bone marrow stromal cells (BMSCs), resulting in FD bone lesions that become apparent during early childhood and lead to deformity, fractures, and pain^(1)^.

FD arises from altered differentiation of bone marrow skeletal stem cells (SSCs), the progenitor subpopulation of BMSCs^(2)^. Affected SSCs are unable to differentiate into either hematopoiesis-supporting marrow stroma or adipocytes and generate instead highly proliferative fibroblastic cells and abnormal osteochondrogenic cells. This leads to regression of normal hematopoietic/fatty marrow tissue into the perimeter of the lesions, and its replacement by fibro-osseous tissue with occasional areas of fibrocartilage. Radiographically, lesions typically appear as lytic/sclerotic “ground glass” areas within the bones, although appearances can vary significantly^(3)^. On histology, they appear as vascularized fibro-osseous tissue of undifferentiated fibroblasts and curvilinear trabeculae of abnormal, poorly mineralized woven bone^(3)^.

FD lesions are often rich in osteoclasts and osteolytic activity, and elevated bone turnover is a characteristic of the disease^(4)^. Indeed, osteoclasts are recruited by an abnormally high production of RANKL by FD BMSCs, to the extent that circulating RANKL levels on FD patients average 16-fold increase that on healthy controls^(5)^. These findings led to the consideration of RANKL inhibition as a therapeutic approach for FD^(6,7)^. A clinical trial performed by our group using the anti-RANKL drug denosumab was shown successfully halt bone resorption in FD, and to prevent proliferation of altered BMSCs, leading to decreased cellularity, and normalized differentiation of affected BMSCs. This resulted in a better mineralized and organized bone in FD lesions and a decrease in co-morbidities associated with the skeletal lesions. Based on these findings, our current understanding of FD pathophysiology is that of a positive feedback loop between altered BMSCs and osteoclasts, through which FD BMSCs release pro-osteoclastic factors inducing osteoclastogenesis, and osteoclasts respond increasing BMSC proliferation and contributing to their altered differentiation through intercellular signals like RANK-containing extracellular vesicles^(8,9)^. Despite this advancement of our understanding of the cellular dynamics in FD tissue, little is known about the transcriptional effects of Gα_s_ activation/cAMP excess in BMSCs and how this translates into their modification of the FD lesion microenvironment through the modulation of released signaling factors.

Therefore, a comprehensive characterization of the genetic expression and production of secreted factors in addition to RANKL and OPG by FD BMSCs is crucial to understand the microenvironment of FD lesions. Moreover, as the discovery of RANKL’s role in FD led to the trial of denosumab therapy, such an exploration would provide novel therapeutic approaches and/or circulating biomarkers for FD. Previous efforts have been done to characterize released factors in FD other than RANKL/OPG, including the demonstration of lesional FGF23 excess^(10)^ and the investigation of IL6 in this disease, that led to the unsuccessful attempt to treat FD patients with the IL6R-inhibiting drug tocilizumab^(11–13)^. However, the scope of these exploratory studies was limited due to the lack of high throughput screening techniques.

On the other hand, we and others carried out efforts to characterize the transcriptomic profile of FD, but our approach involved the isolation of mRNA in bulk FD tissue, which does not allow for determination of the contribution of BMSCs bearing gain-of-function variants of GNAS to the differential expression genes identified ^(9,14–16)^. In addition, previous attempts were done to characterize the transcriptomic changes caused by *GNAS* activation in FD BMSCs using cultured human BMSCs transduced with Gα_s_^R201C^, though the cellular manipulation of this technique may limit its reliability to capture the transcriptomic effects of Gα_s_ gain of function in lesional BMSCs ^(17,18)^.

We developed a methodology which utilized a combination of bulk RNAseq data obtained in cultured BMSCs from an inducible mouse model of FD and in cultured BMSCs of FD patients compared with healthy volunteers. This resulted in the development of a robust transcriptomic signature specific to altered FD BMSCs, including only genes significantly and similarly modulated in both systems. We then used these cultures to measure the concentration of several pro-osteoclastogenic cytokines and other factors of interest secreted into the conditioned culture medium. Lastly, we measured the circulating concentrations of some of these cytokines in plasma samples from FD patients, and correlated them with their disease burden, resulting in the proposal of novel potential circulating biomarkers for FD.

## Materials and Methods

### Human and Mouse Specimens

Primary cultures of bone lesions from six patients with FD, as well as plasma samples from 57 patients with FD were selected from a longstanding natural history protocol (NCT00001727) (Fig 1, Tables 1 and 2). The protocol was approved by the NIH Investigational Review Board, and all subjects/guardians gave informed consent/assent. Subjects were limited to those who never received bisphosphonate or denosumab therapy. All subjects had skeletal disease burden scores (SBS) measured using a previously validated Tc-99m bone scintigraphy method^(19)^. Plasma was collected from patients at NIH and cryopreserved; bone specimens were collected as waste during corrective surgeries and used to culture primary BMSCs, which were subsequently cryopreserved. BMSCs from six healthy donors were obtained according to NIH ethical guidelines (NIH OHSRP exemption #373) (Table 2).

**Figure 1.**
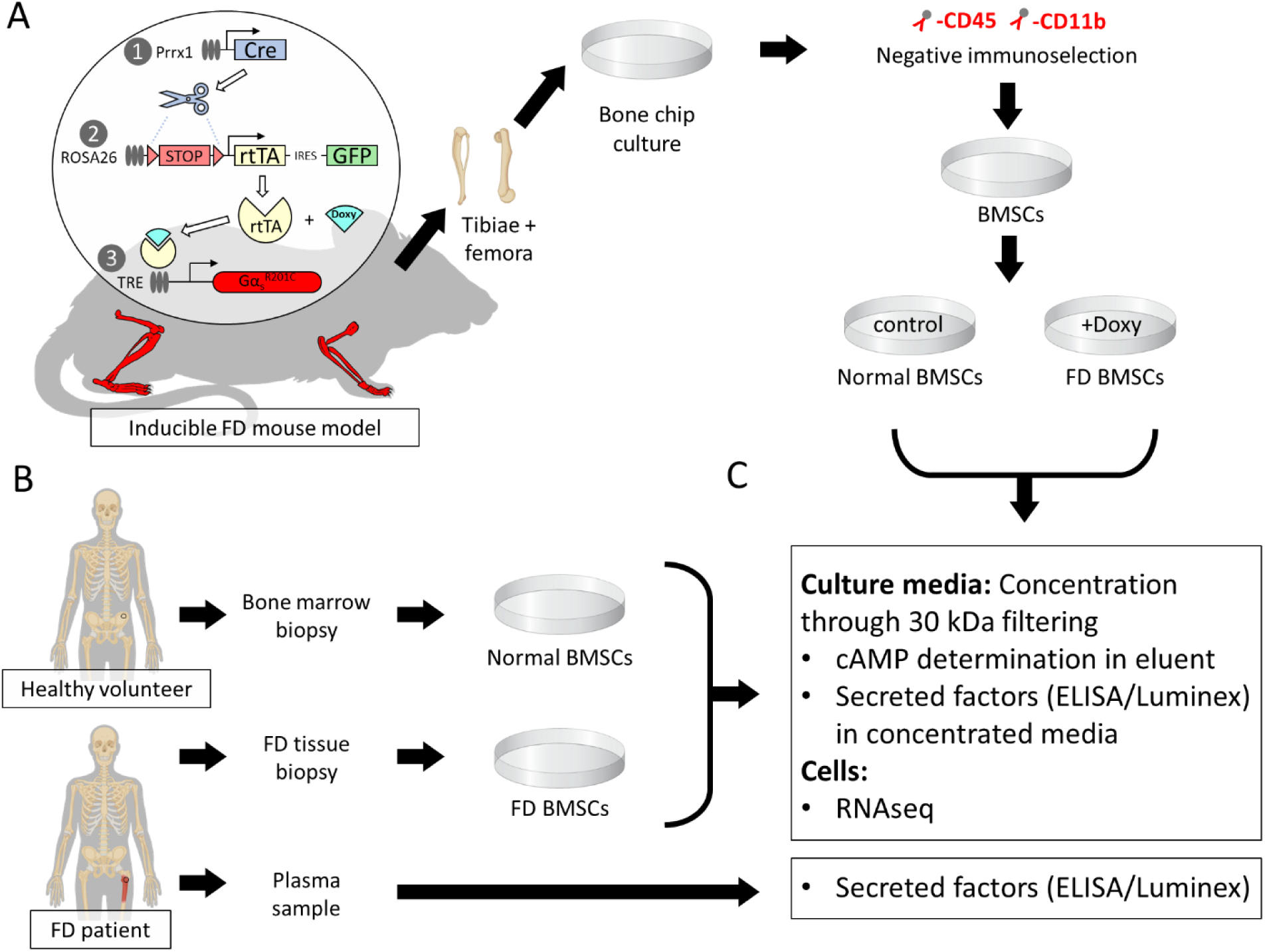
Schematic representation of the study design and workflow. **A)** Briefly, bone marrow stromal cells were cultured and isolated from a mouse model with doxycycline-inducible expression of Gα_s_^R201C^ in cells of the osteogenic lineage in the appendicular skeleton through passage enrichment and negative immunoselection against hematopoietic markers CD45 and CD11b. After that, cells from each mouse were cultured in two groups. At 80% confluency, cultures were transferred to FBS-depleted media and Gα_s_^R201C^ expression was induced in one of the groups by adding 5 mg/mL of doxycycline. **B)** Human BMSCs were cultured from surgical waste FD tissue and bone marrow specimens from healthy volunteers. At 80% confluency, cells were transferred to FBS depleted media. Plasma was obtained from FD patients. **C)** Media was collected from human and mouse cells 48h later, concentrated by filtration, and used to determine cAMP and secreted factor concentrations. Attached cells were lysed with TRIzol for RNAseq. FD patient plasma was used for detecting BMSC-secreted factors in circulation.

**Table I.** FD patient plasma donor demographics.

| **Characteristics (N=57)** | **N (%) or Mean (range)** |
| --- | --- |
| Females | 32 (56%) |
| Age at blood draw (years) | 22 (4-58) |
| Skeletal burden score | 35.46 (0.5-75) |
| MAS features |  |
| Skin hyperpigmentation | 42 (74%) |
| Precocious puberty | 26 (46%) |
| Growth hormone excess | 23 (40%) |
| Hyperthyroidism | 20 (35%) |
| Hypophosphatemia | 23 (40%) |
| Cushing syndrome | 4 (7%) |

**Table II.** BMSC donor demographics.

| **Characteristics** | **FD patients**<br>**N (%) or Mean (range)** | **Healthy volunteers**<br>**N (%) or Mean (range)** |
| --- | --- | --- |
| Females | 3 (50%) | 1 (16.66%) |
| Age at sample donation (years) | 13.33 (3-32) | 27.66 (22-36) |
| Culture heterozygosity (% GNAS p.R201C/H mRNA) | 17.16% (7-55%) | N/A |
| Sample source location |  |  |
| Iliac crest |  | 6 (100%) |
| Femur | 6 (100%) |  |

Femora and tibiae from 4 inducible FD mice carrying a doxycycline-inducible Gα_s_^R201C^ variant were dissected and used to obtain primary murine BMSC cultures (Fig 1). To limit the variability introduced by mixing sexes and ages in the analyses, all mice used were 12-week-old males.

### Human BMSC culture

Cryopreserved BMSCs isolated from 16 patients with FD and 6 healthy volunteers (HV) were cultured as previously described^(20)^ and cryopreserved. Cultures were thawed and seeded in T-25 flasks for continued growth in full medium (α-MEM supplemented with 20% fetal bovine serum and penicillin/streptomycin) until at least 5.5 million cells were present. Cells were detached using trypsin (Sigma-Aldrich, T4049) and the number of total cells was confirmed using a Cellometer Auto M10 (Nexcelom Bioscience). Two T-25 flasks were seeded with 2 million BMSCs in full medium for each cell line; after one day, medium was changed to depleted medium (α-MEM supplemented with penicillin/streptomycin). Cells were then incubated for 48 hours, then culture medium was collected from each well. TRIzol Reagent (Invitrogen, 15596026) was added to flasks and frozen at −80°C. Complete Mini EDTA Free Protease Inhibitor Cocktail (Roche, 4693159001) was added to medium, which was centrifuged to discard aggregates, concentrated 10 times using Amicon Ultra-4 Ultracel-3 3 kDa centrifugal filter units (Millipore, UFC800308). Aliquots of concentrated medium and flowthrough medium were frozen at −80°C.

### Murine BMSC culture

To avoid excessive presence of hematopoietic cells in the primary cultures, mouse bones were depleted of bone marrow, minced into small chips and seeded onto T-25 flasks using full medium (same formulation as in human cultures). Medium was changed after 6 hours and then every 4 days. Bone chips were subsequently removed and placed in new T-25 flasks on days 12 and 21. On day 28, flasks showing high ratio of stromal cells to monocytes by visual examination on an inverted microscope were treated with trypsin (Sigma Aldrich, T4049) and harvested cells were transferred to T-75 flasks. When an estimated 12 million cells per subculture was achieved, cells were trypsinized and passed through a nylon mesh for single cell suspension. The total number of live cells was confirmed using a Cellometer Auto M10 (Nexcelom Bioscience) and Trypan Blue (Sigma-Aldrich, T8154). Cultures were negatively immunoselected for hematopoietic cells using CD45 and CD11b microbeads (Miltenyi Biotec, 130-052-301 & 130-097-142) and the autoMACS Pro Separator following the manufacturer instructions. 6-well plates were seeded with 0.8 million cells per well in full medium. After 48 hours, the medium was changed for depleted medium (same formulation as in human cultures). Half of the cultures from each mouse were treated with doxycycline hyclate (5 mg/mL, Sigma-Aldrich D5207) for 48 hours and media was collected. Complete Mini EDTA Free Protease Inhibitor Cocktail (Roche, 4693159001) was added to media. Cells were then incubated for 48 hours, then culture media was collected from each well. TRIzol Reagent (Invitrogen, 15596026) was added to flasks and frozen at −80°C. Media was centrifuged to discard aggregates, concentrated 10 times using Amicon Ultra-4 Ultracel-3 3 kDa centrifugal filter units (Millipore, UFC800308). Aliquots of concentrated media and flowthrough media were frozen at −80°C.

### RNA Extraction, Sequencing, and Analysis

Bulk RNA was extracted from cultured cells using the TRIzol Reagent protocol without glycogen. Library preparation and sequencing was performed by the Genomics and Computational Biology Core (NIDCR, NIH) using an Illumina HiSeq 2500 (Illumina, San Diego, CA) configured for 37 paired-end reads (human) or 150 paired-end reads (mouse). Read quality was assessed using FastQC software^(21)^ and reads were subsequently mapped using STAR aligner v2.5.2a ^(22)^. Mapping quality was assessed using Picard tools. Read counts were determined using the quantMode utility of STAR aligner and genes with less than 5 counts in at least 1 sample were filtered out. Normalized counts were then calculated using TMM normalization. Cultures showing GNAS p.R201C/H mRNA variant allele expression frequency of 7-55% were selected for further analyses (6 out of 16 samples cultured). Briefly, bam alignments were visualized using the Integrative Genome Viewer (IGV) application and the nucleotide counts at the GNAS p.R201 position and corresponding resulting aminoacid changes were determined for each sample. ^(23)^. DEseq2^(24)^ was used to determine differential gene expression. Unsupervised clustering was performed using principal component analysis and unsupervised clustering using heatmap.2 in R^(25)^. Other heatmaps were manually generated by grouping related genes that were differentially regulated in the same direction.

Cross-species human-mouse annotation was performed using the Biomart service using the high-confidence ortholog dataset. Annotated genes were then used to determine the number of genes differentially regulated in the same manner between both species (defined as the FD Signature). Pathway analysis was performed for human, mouse, and the FD signature using Enrichr (https://maayanlab.cloud/Enrichr/) to generate bar charts from GO Molecular Function 2021, BioPlanet 2019, KEGG 2021 Human, and MSigDB Hallmark 2020 using |log_2_-fold change| > 1.5 and adjusted p-value < 0.01.

Predicted protein-protein interactions were mapped based on genes differentially regulated in the same direction in both species (adjusted p < 0.01) using the STRING database (v11.5; string-db.org). The top 6 groups resulting from MCL clustering were annotated based on common protein functions.

### cAMP Determination

cAMP-Gs HiRange standard (Cisbio, 62AM6CDA) was dissolved in 11.2 mM 3-isobutyl-1methylxanthine (IBMX; Sigma-Aldrich, I5879 and Dulbecco’s Modified Eagle Medium (DMEM; Gibco 10564011) for external calibration. A 200 µL aliquot of post-filtration flowthrough media was combined with 500 µM IBMX to protect from cAMP degradation. The aliquot was combined with 200 µL acetonitrile and centrifuged (13,0000 G, 30 min); 50 µL of supernatant was combined with 50 µL HPLC-grade water.

Liquid chromatography with tandem mass spectrometry was then used to quantify cAMP using an orbitrap Fusion Lumos mass spectrometer interfaced to an Ultimate3000 HPLC system (Thermo Scientific, West Palm Beach, FL) and an Atlantis C18 column (2.1×150mm, 3µm, Waters Corp, Milford, MA). Samples were injected and desalted for 2 min at 2% solvent B (80% acetonitrile, 20% water, and 0.1% formic acid) in solvent A (0.1% formic acid in water), followed by a linear gradient to 98% B in 3 min, the composition was held at 98% B for 2.5 min before ramping down to 2% B in 0.1min. Column was equilibrated for 4.5 min at 2% B before the next injection. Flow rate was 250 µL/min. Solvent A and solvent B were Optima grade from Fisher Scientific. Parallel Reaction Monitoring of protonated cAMP at m/z 330.0598 ([M+H]+) was recorded in the orbitrap at R=50,000 for detection and quantification using positive ion electrospray ionization with the following source parameters: Spray voltage 3300V, Sheath gas 40 unit, Aux gas 10 units, Sweep gas 2 units. The ion transfer tube was at 325 °C, and vaporizer at 350 °C. Isolation window was 1.2 m/z, collision energy was 35%. Standard curves were acquired before and after the sample set. A QC sample (5nM standard) was analyzed mid-sequence. Quantification was carried out using QuanBrowser module of Xcalibur software (Thermo Scientific) using peak area of m/z 136.0609. Quantification result was manually inspected before exporting to Excel.

### Serum and Media Determinations

For mouse samples, RANKL was detected using Mouse TRANCE/RANKL/TNFSF11 Quantikine ELISA Kit (R&D systems, MTR00), Ephrin B2, Semaphorin 3A and FAPα using ELISAs from LSBio (LS-F6974, LS-F33608 and LS-F49804, respectively) and soluble Fas ligand and M-CSF using R&D Quantikine assays (MFL00 and MMC00B, respectively). In addition, a customized magnetic Luminex assay was designed to assess Dkk-1, IL-7, MMP-2, β-NGF, and VEGF in mice (R&D Sytems, LXSAMSM-05 with BR77, BR14, BR37, BR43, and BR21). Mouse cytokines were detected with the Bio-Plex Pro Mouse Cytokine 23-plex Assay (Bio-Rad, #M600009RDPD), including, IL-1α, IL-1β, IL-2, IL-3, IL-4, IL-5, IL-6, IL-9, IL-10, IL-12 (p40), IL-12 (p70), IL-13, IL-17A, Eotaxin, G-CSF, GM-CSF, IFN-γ, KC, MCP-1 (MCAF), MIP-1α, MIP-1β, RANTES, and TNF-α.

In human samples, RANKL was detected using Biomedica Immunoassay for free sRANKL ELISA kit (Eagle Biosciences, BI-20462), and cytokines were detected using the Bio-Plex Pro Human Cytokine 17-plex Assay (Bio-Rad, #M5000031YV), including IL-1β, IL-2, IL-4, IL-5, IL-6, IL-7, IL-8, IL-10, IL-12 (p70), IL-13, IL-17A, G-CSF, GM-CSF, IFN-γ, MCP-1 (MCAF), MIP-1β, TNF-α.

Analytes that were undetectable in human or mouse experimental and control groups were excluded from analyses.

### Statistical Analysis

Comparisons between released cytokines in cell culture were performed on GraphPad Prism v9.2.0 (GraphPad Software, Boston, MA) and determined using Multiple Mann-Whitney tests with a two-stage Benjamini, Krieger, and Yekutieli step-up and 5% False Discovery Rate (FDR). The correlations between circulating cytokines and skeletal burden score in patients with FD/MAS were determined using multiple correlations with Pearson r in GraphPad. Using SAS v9.4 (SAS Institute, Cary, NC), p-values in SBS-cytokine and cytokine-cytokine correlations were adjusted for multiple testing, with an FDR-adjusted p-value below 0.05 considered statistically significant. Unpaired t-tests were used to determine significance between cAMP and variant burden comparisons with a threshold of p<0.05. Statistical analysis of differential gene expression was included in the DESeq2 package.

## Results

### Murine and human Gα_s_^R201C/H^-expressing BMSC cultures display pro-inflammatory transcriptomic profiles

Bone chips from both femora and tibiae from each mouse were minced and plated. Two to four weeks later, cultures were depleted of CD45+ CD11b+ hematopoietic cells and seeded in 6-well plates. Two days later, half of the cultures were induced for Gα_s_^R201C^ expression for 48h by supplementation of the culture media with doxycycline (5 mg/mL), and mRNA was extracted (Fig 1). Gα_s_^R201C^ transgene expression was confirmed in the transcriptomic analysis (Fig 2A) as well as downstream pathway activation by measurement of cAMP in the culture media (Fig 2B). RNAseq transcriptomic analysis showed good segregation by Gα_s_^R201C^ expression status based on principal component (PC) analysis, which also grouped together on unsupervised clustering (Fig 2C, D) and revealed significant modulation of 5,852 genes (Table S1). Pathway analysis of the most differentially expressed genes in this dataset (with at least |log2 fold change| > 1.5 and adjusted p<0.01) was performed using Gene Ontology (GO) Molecular Function, BioPlanet, Kyoto Encyclopedia of Genes and Genomes (KEGG), and Molecular Signatures Database (MSigDB) hallmark gene set repositories with the Enrichr tool. Some of these gene sets showed signatures directly or indirectly associated to Gα_s_ activation, and all of them showed terms related to cytokine signaling and inflammation: “cytokine activity” in GO Molecular Function; “TGF-beta regulation of extracellular matrix” and “Interleukin-1 regulation of extracellular matrix” in BioPlanet; “Cytokine-cytokine receptor interaction” in KEGG; “TNF-alpha Signaling via NF-kB” and “Inflammatory Response” in MSigDB (Fig 2E).

**Figure 2.**
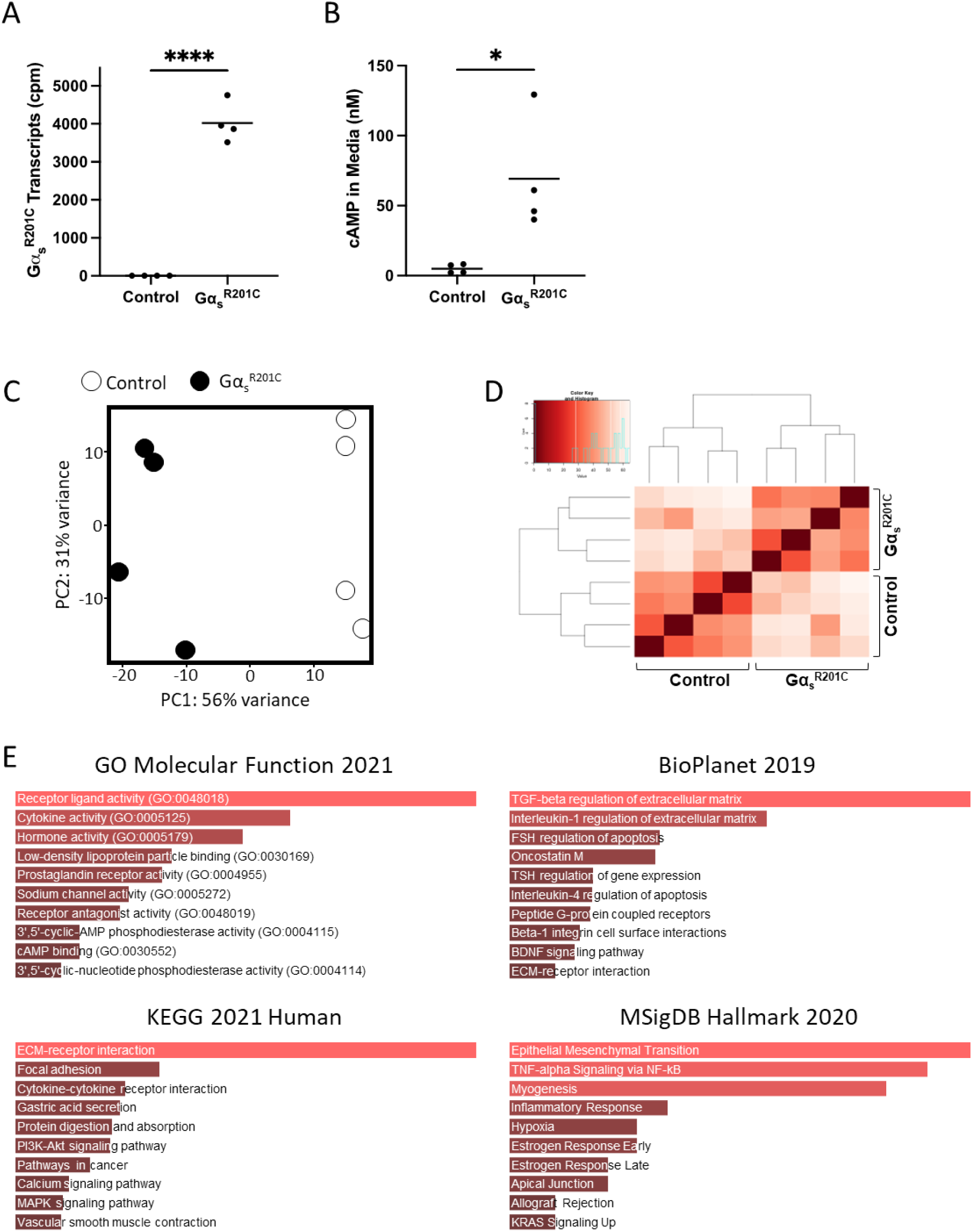
Pro-inflammatory pathways are enriched in BMSCs derived from an inducible mouse model of FD. **A)** Transcriptomic data confirm that Gα_s_^R201C^ is highly expressed by murine BMSCs following induction with doxycycline. **B)** BMSCs expressing Gα_s_^R201C^ exhibited markedly elevated cAMP concentrations in culture media, as determined by HPLC-MS. **C, D)** Principal component and unsupervised clustering analyses demonstrated close grouping within control and induced experimental groups while maintaining segregation between groups, confirming robust differences in expression profiles. **E)** Gene set enrichment analyses of significantly regulated genes (|log_2_FC| > 1.5 and p-adj < 0.01) revealed enriched pathways for inflammatory signaling, e.g., “cytokine activity” (GO Molecular Function 2021); “TGF-beta regulation of extracellular matrix” and “Interleukin-1 regulation of extracellular matrix” (BioPlanet 2019); “cytokine-cytokine receptor interaction” (KEGG 2021 Human); and “TNF-alpha Signaling via NF-kB” (MSigDB Hallmark 2020). *p<0.05, ****p<0.0001

Of all 16 human FD BMSCs cultures tested, six had an Gα_s_^R201C/H^ mRNA allele frequency 7-55% of all Gα_s_ transcripts reads and were selected for further analyses and compared to BMSCs obtained from healthy volunteers (Fig 3A). Gα_s_ activation was confirmed through measurement of cAMP in the culture media (Fig 3B). PC analysis showed excellent segregation of FD and healthy volunteer (HV) BMSCs cultures, and both categories partially grouped together in unsupervised clustering analysis (Fig 3C, D) and revealed significant modulation of 3,637 genes (Table S2). We performed a similar Enrichr pathway analysis as in mouse cultures, with the same fold change and p-value restrictions, which also showed pathways consistent with Gα_s_ activation, and all showed gene sets related to cytokines and inflammation: “cytokine activity” and “chemoattractant activity” in GO Molecular Function; “TGF-beta regulation of extracellular matrix” and “Cytokine-cytokine receptor interaction”, “Interleukin-4 regulation of apoptosis”, “TGF-beta regulation of skeletal system development” in BioPlanet; “Cytokine-cytokine receptor interaction”, “TGF-beta signaling pathway” in KEGG; “TNF-alpha Signaling via NF-kB” and “Inflammatory Response” in MSigDB (Fig 3E).

**Figure 3.**
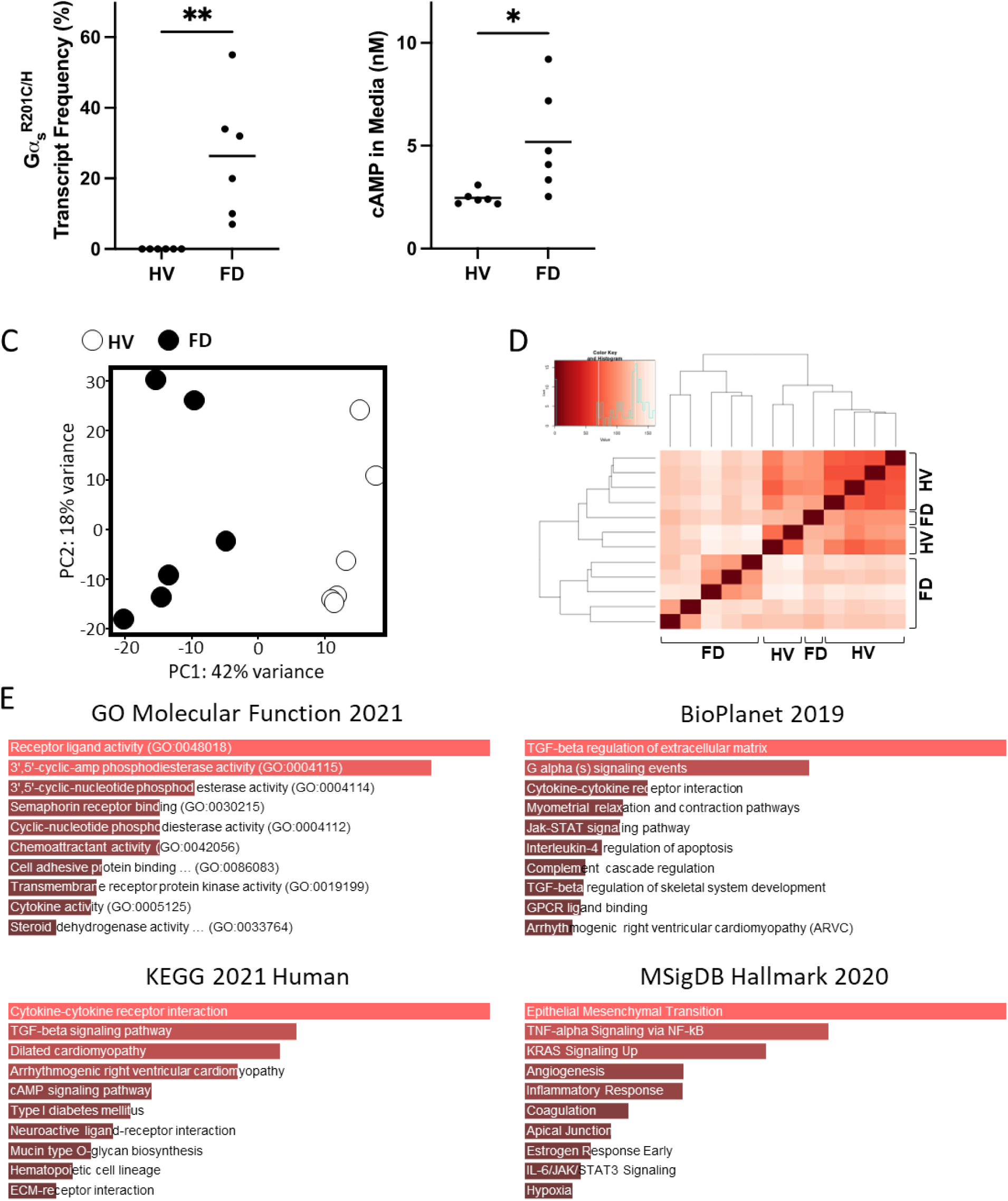
Primary BMSCs from patients with FD also exhibit a pro-inflammatory phenotype. **A)** Transcriptomic data confirmed expression of Gα_s_ variants in BMSCs from patients with FD. Notably, the fractional abundance of variant transcripts was highly variable, reflecting the mosaic nature of the disease. **B)** HPLC-MS demonstrated elevated cAMP production in FD cultures compared to healthy volunteers (HVs), in line with Gα_s_ activation. **C, D)** Principal component and unsupervised clustering analyses showed close grouping of HV and FD samples while maintaining segregation between groups. **E)** Gene set enrichment analyses of significantly regulated genes (|log_2_FC| > 1.5 and p-adj < 0.01) revealed enriched pathways for inflammatory signaling, e.g., “cytokine activity” and “chemoattractant activity” (GO Molecular Function 2021); “TGF-beta regulation of extracellular matrix,” “Interleukin-4 regulation of apoptosis,” and “TGF-beta regulation of skeletal system development” (BioPlanet 2019); “cytokine-cytokine receptor activity” (KEGG 2021 Human); and “TNF-alpha Signaling via NF-kB” (MSigDB Hallmark 2020). *p<0.05, **p<0.01

### Combining the transcriptome of human and murine FD BMSCs reveals a robust genetic signature

To develop a robust signature of genes differentially modulated in FD BMSCs across disease models, we combined the datasets of differentially expressed human and mouse genes in wildtype Gα_s_ and Gα_s_^R201C/H^-expressing BMSCs. Of the 3,367 differentially expressed genes in human samples, 2,964 had a mouse ortholog; and of the 5,852 differentially expressed mouse genes, 5,355 had a human ortholog. Of these, 1,239 genes appeared in both datasets and 764 were regulated in the same direction (Fig 4A, Table S3). A selection of these genes involved in pathways of interest for the study of FD curated by literature review are shown in Figure 4B, including genes involved in Gα_s_/cAMP signaling, SSC differentiation, fibrosis, inflammation (including pro-osteoclastogenic cytokines), and vascularization. In addition, the 136 genes with strongest regulation (defined by |log_2_ fold change| > 1.5 and adjusted p < 0.01) were analyzed with Enrichr pathway analysis, showing over-representation of genes related to cAMP signaling in GO Molecular Function; “G alpha s pathway” in BioPlanet; “Cytokine-cytokine receptor interaction” in KEGG; and “TNF-alpha Signaling via NF-kB” and “Inflammatory Response” in MSigDB (Fig 4C). Lastly, we selected those genes with a p < 0.01 (435 genes) to assess the pathway interaction of their encoded proteins using STRING, which detected 6 main interacting pathways, involved in processes like cAMP signaling, extracellular matrix organization, metallopeptidase activity, cellular metabolism, mesenchymal cell proliferation, and ATPase activity (Fig S1).

**Figure 4.**
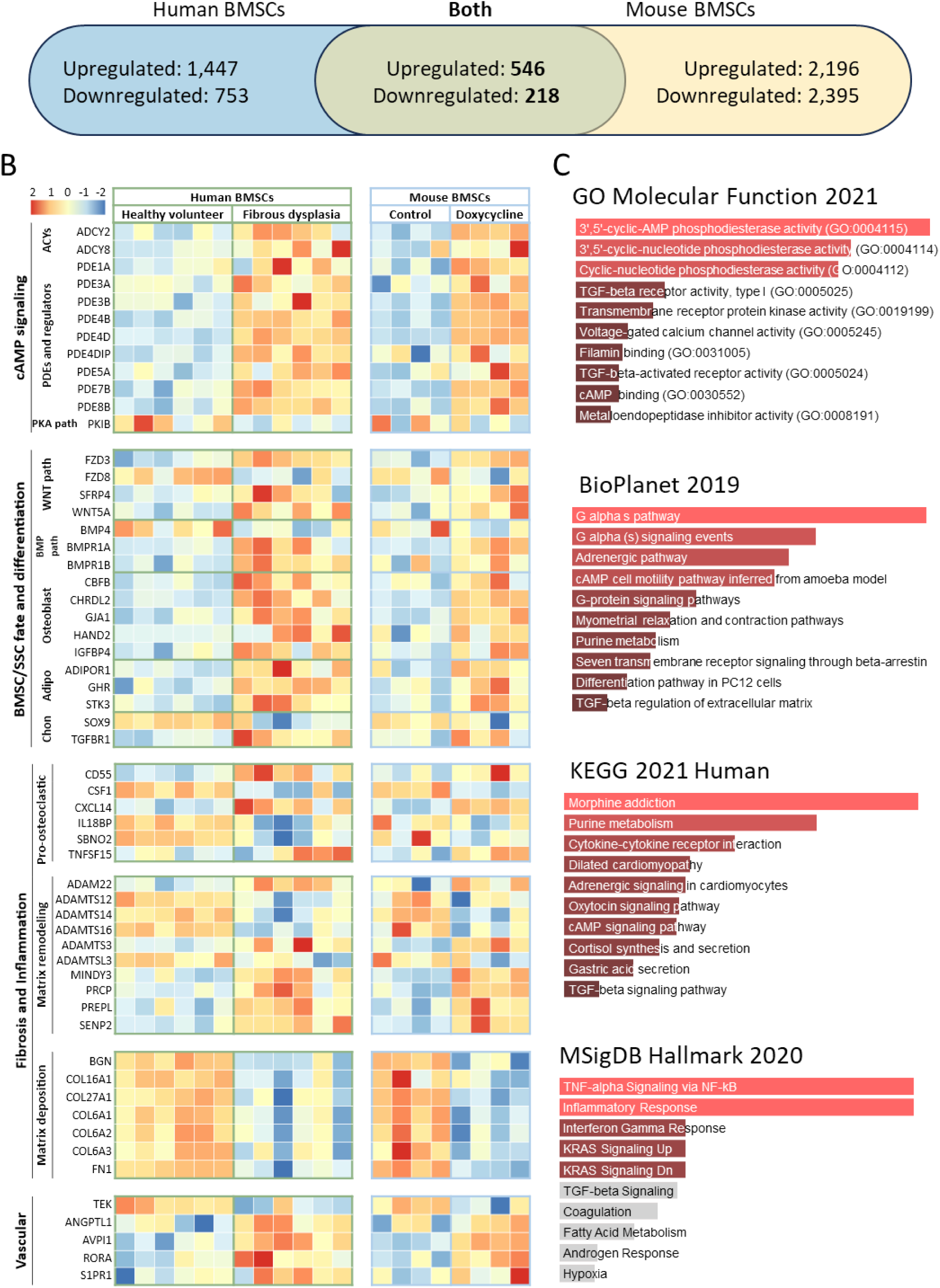
BMSCs from mice and humans with FD shared genes dysregulated in the same manner. **A)** Differentially expressed genes (p-adj < 0.05) were limited to those having orthologs in either species, resulting in 2,964 and 5,355 differentially expressed human and mouse genes, respectively. Of these, 764 genes appeared in both species and were differentially expressed in the same direction. **B)** From our FD Signature, genes were manually selected and tabulated according to known associations, e.g., with the WNT and BMP pathways. **C)** Gene set enrichment analysis of our FD Signature revealed enriched pathways similar to previous results from mice and humans, e.g., “cytokine-cytokine receptor interaction” (KEGG 2021 Human) and “TNF-alpha Signaling via NF-kB” (MSigDB Hallmark 2020). ACYs, adenylyl cyclases; adipo, adipogenesis; chon, chondrogenesis; PDEs, phosphodiesterases; SSC, skeletal stem cell.

### Mouse Gα_s_^R201C^-expressing BMSC cultures release pro-inflammatory cytokines and other factors related to FD pathophysiology

We measured the level of 18 cytokines in the primary culture media of BMSCs from FD patients and healthy volunteers (HV). Of these, five cytokines were undetectable and due to the high variability of the detected levels in these samples we failed to detect significantly different levels of any of the 12 remaining cytokines between HV and FD BMSC culture media (Fig S2). However, we measured 24 cytokines in media from murine BMSC cultures expressing or not Gα_s_^R201C^, of which 13 factors were significantly higher in Gα_s_^R201C^-expressing BMSCs (Fig 5), nine did not change (Fig S3), and two fell below the detection range of the assays (IL4 and IL9). In addition to these cytokines, we measured other factors involved in the FD microenvironment, such as the proteases MMP2 and FAPα, the WNT modulator Dkk1, the growth factors VEGF and β-NGF and modulators of osteoclastogenesis previously shown to be produced by osteogenic cells (EPHD4, FASL and SEMA3A). Although the assays used could not detect presence of EFNB2 and FASL in the media, all the remaining factors showed increased levels in cultures with induced Gα_s_^R201C^ expression (Fig 6).

**Figure 5.**
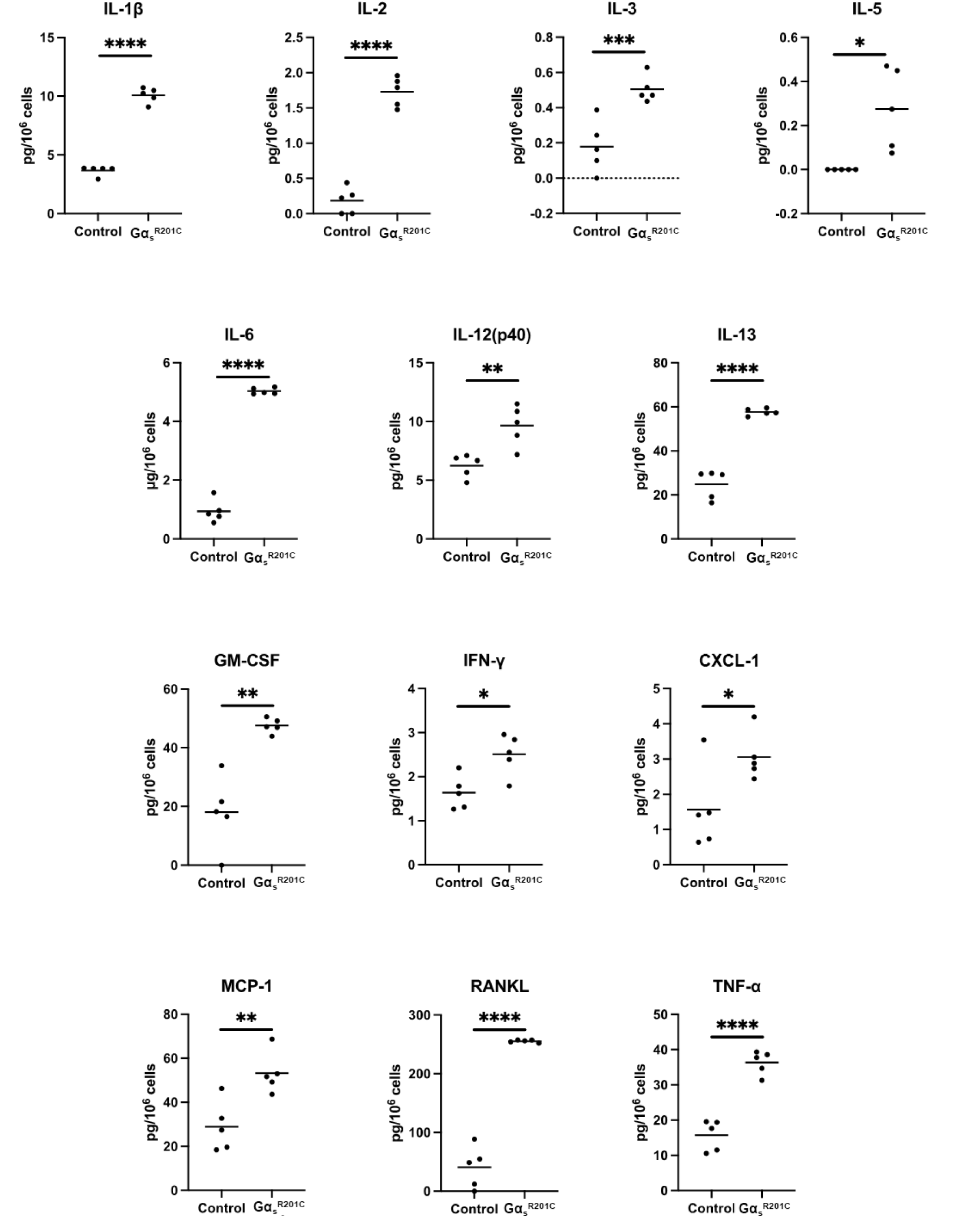
Cultured BMSCs from FD mice release pro-inflammatory cytokines. Multiple Mann-Whitney test with two-stage Benjamini, Krieger, and Yekutieli and 5% FDR was used to determine statistical significance. IL-4 and IL-9 were undetectable. *p<0.05, **p<0.01, ***p<0.001, ****p<0.0001.

**Figure 6.**
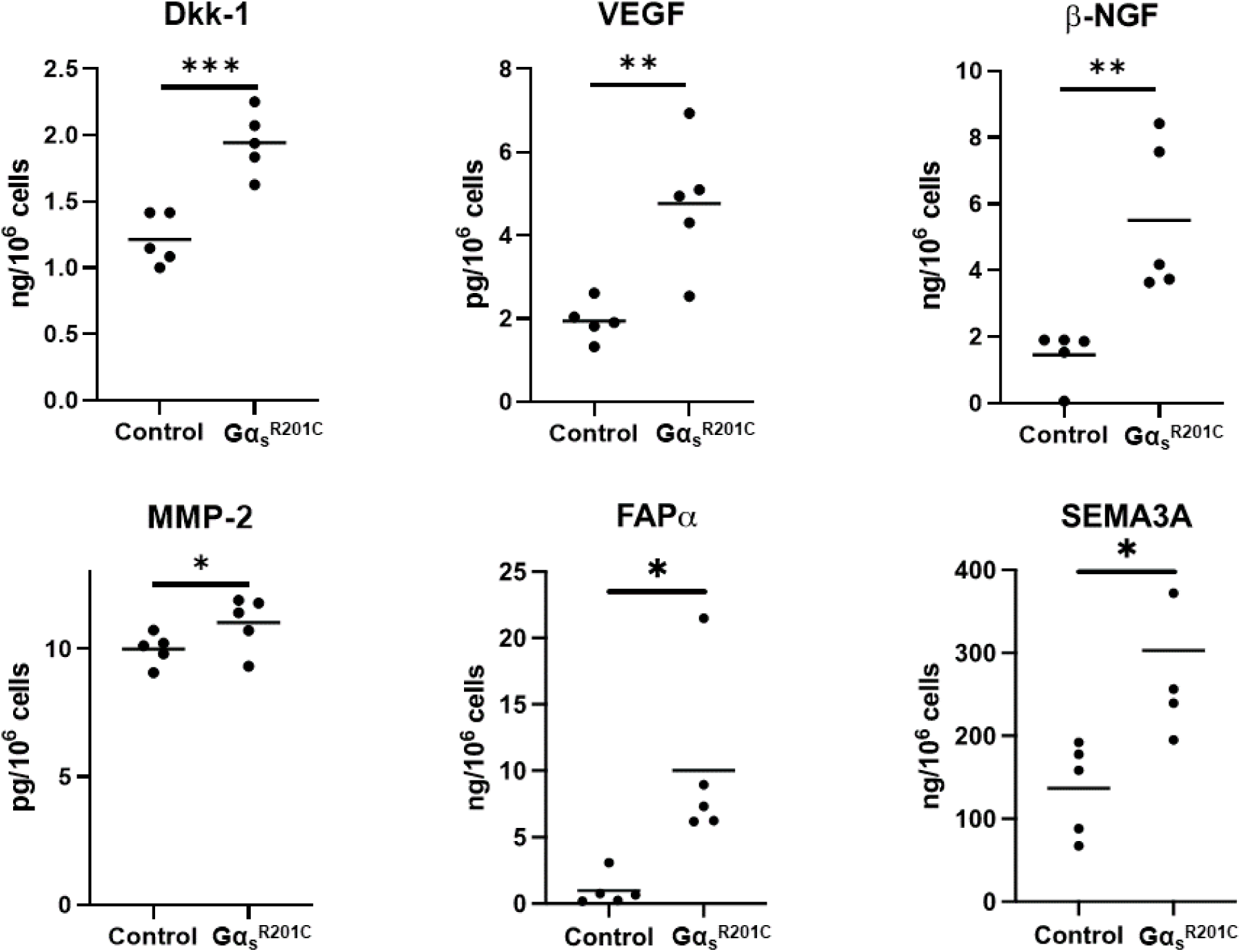
Cultured BMSCs from FD mice release additional factors related to FD pathogenesis. Multiple Mann-Whitney test with two-stage Benjamini, Krieger, and Yekutieli and 5% FDR was used to determine statistical significance. Ephrin B2 and soluble Fas ligand were assessed but not found in detectable levels. *p<0.05.

### Plasmatic cytokines in FD patients correlate with their disease burden

We analyzed 19 cytokines, of which 16 were detectable, in serum samples from 57 patients with FD, with disease burden ranging from a skeletal burden score (SBS) of 0.5 to 75 (Table 1). In addition, the bone turnover markers ALP, osteocalcin and NTX were measured. As expected, ALP, osteocalcin, OPG and RANKL significantly correlated with patient’s SBS. But in addition to these, and for the first time, we demonstrated positive correlation of 7/16 cytokines to SBS (Fig 7A). Interestingly, several cytokines correlated with one another and with ALP and osteocalcin, especially all interleukins analyzed, which presented very strong correlations ranging between r=0.556, p= 2.7*10^-6^ (IL-6 vs IL-7) and r=0.881, p= 3.6*10^-21^ (IL-7 vs IL-4) (Fig 7B).

**Figure 7.**
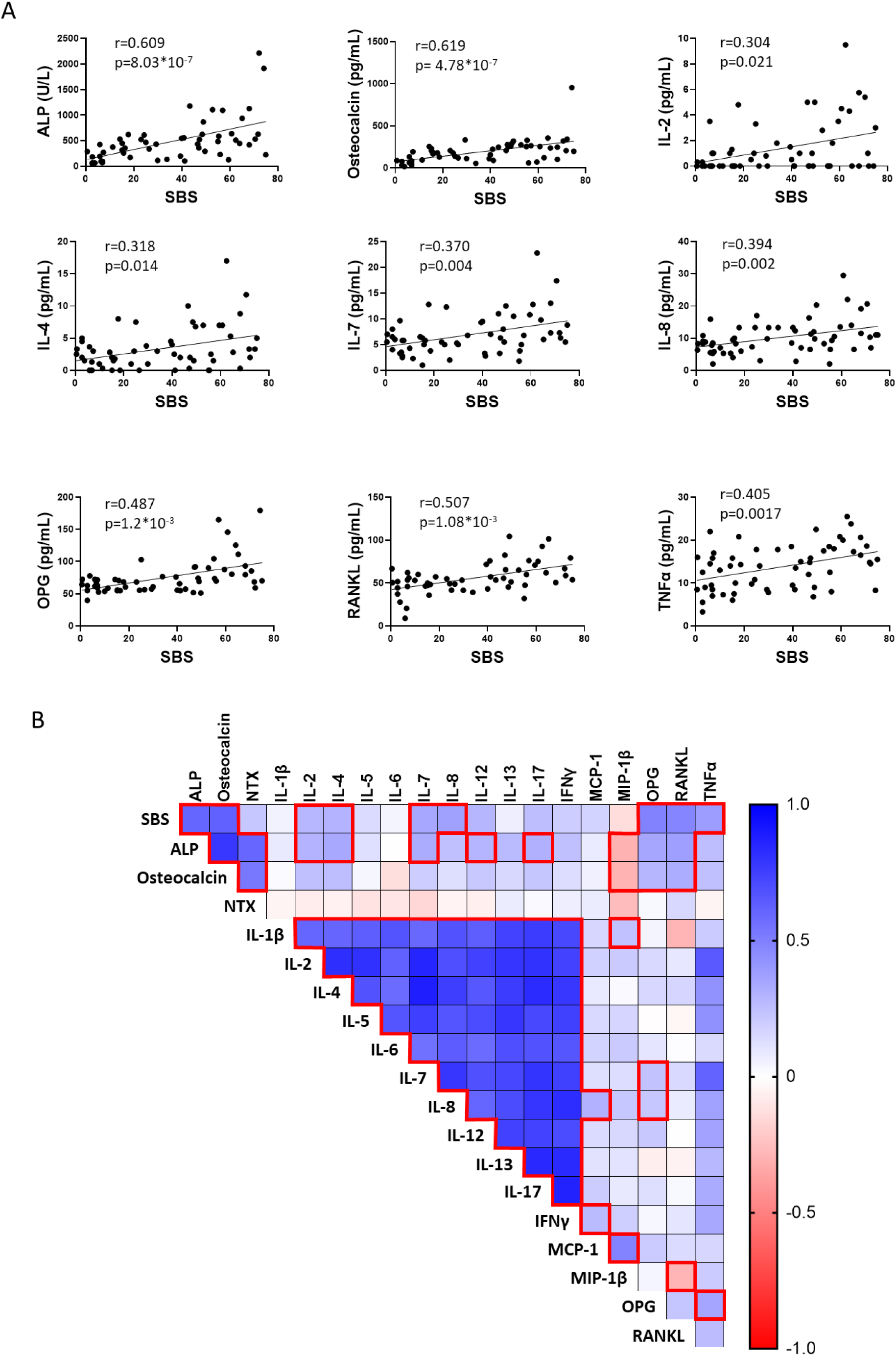
Inflammatory cytokines and bone turnover markers can be detected in serum from patients with FD and correlate with disease burden. **A)** Cytokines and other markers were analyzed in serum from 57 patients with FD and correlated with Skeletal Burden Score (SBS), a standardized quantification of disease burden. Significance was determined by multiple correlations with Pearson r and defined as p<0.05. **B)** SBS-cytokine and cytokine-cytokine correlations between all detectable factors revealed numerous associations, especially with ALP and osteocalcin, as well as between interleukins as expected (red boxes). P-values were adjusted for multiple testing with an adjusted p-value less than 0.05 considered statistically significant. IL-10, G-CSF, and GM-CSF were undetectable.

## Discussion

Although our understanding of the lesional cell population dynamics in FD pathogenesis has advanced significantly in recent years, we still have limited knowledge of the transcriptional effects of hyperactive Gα_s_ and cAMP excess in BMSCs, the underlying cause of the lesions. Previous differential gene expression analyses of bulk FD tissue fail to capture FD BMSCs-specific transcriptomic changes ^(9,14,15)^, and attempts seeking to characterize the effects of Gα_s_^R201C^ expression in human BMSCs by lentiviral transduction of wildtype BMSCs involved high cell manipulation, limiting the model’s validity to emulate the transcriptomic profile of FD BMSCs^(17,18)^. In the present study, to analyze the cell-intrinsic effects of *GNAS* gain-of-function in FD BMSCs unaffected by the lesional microenvironment, we performed a comprehensive exploration of the transcriptome and secretome of FD BMSCs cultured in the absence of other cell types. For a robust experimental design, we combined human and mouse FD cultures to propose a FD BMSC transcriptomic signature, comprised by significantly modulated orthologous genes changing in the same direction in both models.

For both models, *GNAS* variant expression and cAMP release in the culture media were confirmed before data analysis. Principal component and unsupervised clustering analyses of human and mouse cultures showed good segregation of the samples by group (WT/ Gα_s_^R201C^ in mice, HV/FD in humans) and Enrichr analysis identified several terms associated to GPCR/Gα_s_/AC pathway activation, supporting the experimental validity of our approach. In addition, Enrichr analyses of the human or mouse gene lists, as well as the combined FD BMSC signature, undoubtedly supported the importance of cytokine signaling in FD pathogenesis, returning terms like “cytokine activity”, “cytokine-cytokine receptor interaction” or “inflammatory response” among the top matching genetic datasets. Further analysis of the FD BMSCs signature with the STRING protein-protein network analysis tool also confirmed cAMP signaling activation and identified additional expected processes in FD pathogenesis, like “extracellular matrix organization”, “metallopeptidase activity” and “mesenchymal cell proliferation” among others. Lastly, we curated a subset of 57 genes among the 764 genes identified in the FD Signature with biological significance in the disease (Fig 3B) and demonstrated relationship with cAMP signaling, SSC differentiation fate, osteoclast recruitment, fibrous matrix deposition/remodeling and vascularization. This restrictive strategy to elaborate an FD BMSC signature from human and murine cultures likely missed the identification of some genes involved in FD BMSC pathogenesis, due for example to the absence of known gene orthologs between species (10-12% of the genes modulated in each dataset lacked an identified ortholog in the other species). However, at this expense of losing sensitivity, our analysis gained specificity, because the 764 genes identified were regulated by GNAS gain of function in BMSC across models, avoiding model-specific biases or artifacts. Although human lesional BMSC cultures obviously best represent lesional BMSC transcriptome, the need to use healthy volunteer BMSC control cultures to compare gene expression introduces several biases. First, BMSCs were obtained from different skeletal sites and by a different technique (iliac crest aspirates in HV versus femur surgical waste tissue in FD patients). In addition, most (4/6) of the patients with FD that donated surgical tissue for culture were children, which cannot be age-matched to healthy volunteer donors. Last, Gα_s_ ^R201C/H^ mRNA allele frequency in FD BMSC cultures was very variable (7-55%). This biological variability translated to the results, as reflected by our inability to show significant differences in the release of the secreted factors studied between FD and HV BMSCs. On the other hand, mouse-derived BMSC cultures impeccably account for these sources of variability, as paired comparisons can be done between cultures from the same donor, induced or not in vitro to express Gα_s_ ^R201C^. However, they are a less faithful representation of FD BMSCs, as Gα_s_^R201C^ transgene expression is not driven by *GNAS* endogenous promoter, and expression is induced for 48h, lacking the long-term epigenetic effects of constitutive activation of Gα_s_ across several cell generations. To build upon our findings, future studies involving techniques like spatial transcriptomics and single cell RNA sequencing may constitute useful approaches to study FD BSMCs transcriptomics.

Our transcriptomic analysis highlighted the important influence of *GNAS* variant-bearing BMSCs in the local microenvironment of FD lesions, ultimately leading to their characteristic hallmarks. Indeed, as patients age, the number of BMSCs bearing *GNAS* variants decreases, and the cellular composition of FD lesions normalizes^(26)^. We and others previously pointed to the importance of the cytokine RANKL in fibrous dysplasia^(5)^, and this lead to the development of targeted therapies, first in preclinical and compassionate treatment case studies ^(6,7,27–29)^ and then in a clinical trial^(8,9)^, including ongoing studies (https://clinicaltrials.gov/study/NCT05419050). While early studies on the role of RANKL signaling in FD facilitated this promising therapeutic approach, little exploration has been done to find additional cytokines and factors secreted by FD BMSCs that may affect osteoclastic-osteoprogenitor crosstalk, as well as other important features of the FD pathogenesis, such as fibrous tissue deposition and remodeling, angiogenesis, and nociception. Therefore, we carried out a systematic exploration of cytokines and other selected secreted factors in our culture models.

Due to the biological variability of human FD and HV BMSC cultures, we failed to identify significant changes in their media concentrations of the 18 cytokines analyzed. However, measurements in mouse cultures media captured a generally pro-osteoclastic, pro-inflammatory secretome, showing increased levels in 13 of the 25 cytokines examined. RANKL and IL-6 had already been associated to FD pathogenesis ^(5,30)^, although a clinical trial targeting IL-6 pathway failed to produce significant disease improvement^(13)^. Previous analyses failed to detect IL-1α and IL-1β, key pro-inflammatory cytokines, in human FD BMSC cultures^(11)^ but both were detectable in the FD mouse cultures, with IL-1β concentration significantly increased in Gα_s_^R201C^-expressing BMSCs. Another key cytokine in the inflammatory response, TNF-α, was dramatically higher in media from Gα_s_^R201C^-expressing BMSC cultures. We then analyzed the concentration of 18 cytokines as well as bone turnover markers known to correlate with disease burden in a collection of plasma from patients with FD. Six cytokines showed correlation to disease burden, and additional correlations of cytokines to bone turnover markers and to one another were identified, confirming the role of pro-inflammatory cytokines in FD pathogenesis.

Six additional secreted factors were identified in the media of mouse FD BMSCs: Dkk-1 is a Wnt inhibitor that has been shown to promote osteoclast activity and prevent osteoblast differentiation in response to TNF-α^(31)^. VEGF, an important pro-angiogenic factor, is also up-regulated by TNF-α, and has anabolic effects both in osteoclasts and osteogenic cells^(32)^. β-NGF, a neurotrophic factor that is also sensitive to pro-inflammatory microenvironments, promotes axonal outgrowth in nearby nociceptive neurons leading to pain^(33)^. SEMA3A has been proposed as well as a major mediator of nerve regulation of bone turnover as well as a crosstalk factor between osteoblasts and osteoclasts^(34,35)^. Last, MMP-2 and FAP-α are proteases involved in fibrotic tissue remodeling whose mRNA levels were found significantly upregulated in previous studies and may be useful as circulating biomarkers or even targetable factors for FD therapy^(9,15,18,36–40)^.

In conclusion, our study provided a comprehensive analysis of the transcriptomic signature and relevant secreted factors of abnormal BMSCs in FD. By combining data from human and mouse FD models, we identified a robust BMSC FD genetic signature, supporting association of Gα_s_ pathway activation in these cells with cytokine signaling and extracellular matrix reorganization. Our FD secretome profiling identified several pro-inflammatory, pro-osteoclastic cytokines that can be detected in circulation in association with disease severity, as well as other factors relevant in FD pathogenic processes like vascularization, fibrosis, osteoblast-osteoclast crosstalk, and nociception. These findings open novel diagnostic and therapeutic avenues that may be explored in future studies.

## Acknowledgements

This research was supported by the Division of Intramural Research, National Institute of Dental and Craniofacial Research, a part of the Intramural Research Program of the National Institutes of Health, Department of Health and Human Services. This research was also made possible through the NIH Medical Research Scholars Program, a public-private partnership supported jointly by the NIH and contributions to the Foundation for the NIH from the Doris Duke Charitable Foundation, Genentech, the American Association for Dental Research, the Colgate-Palmolive Company, and other private donors. We thank Xiaobai Li for her excellent advice and work in statistical analysis of this data, and to Rebeca Galisteo for her continuous support in the laboratory. We also appreciate the dedication of the personnel of NIDCR Veterinary Resources Core to the care of the animals in this study and the guidance in the RNAseq analyses by NIDCR Genomics and Computational Biology Core.

## Supplementary Figures

**Figure S1.**
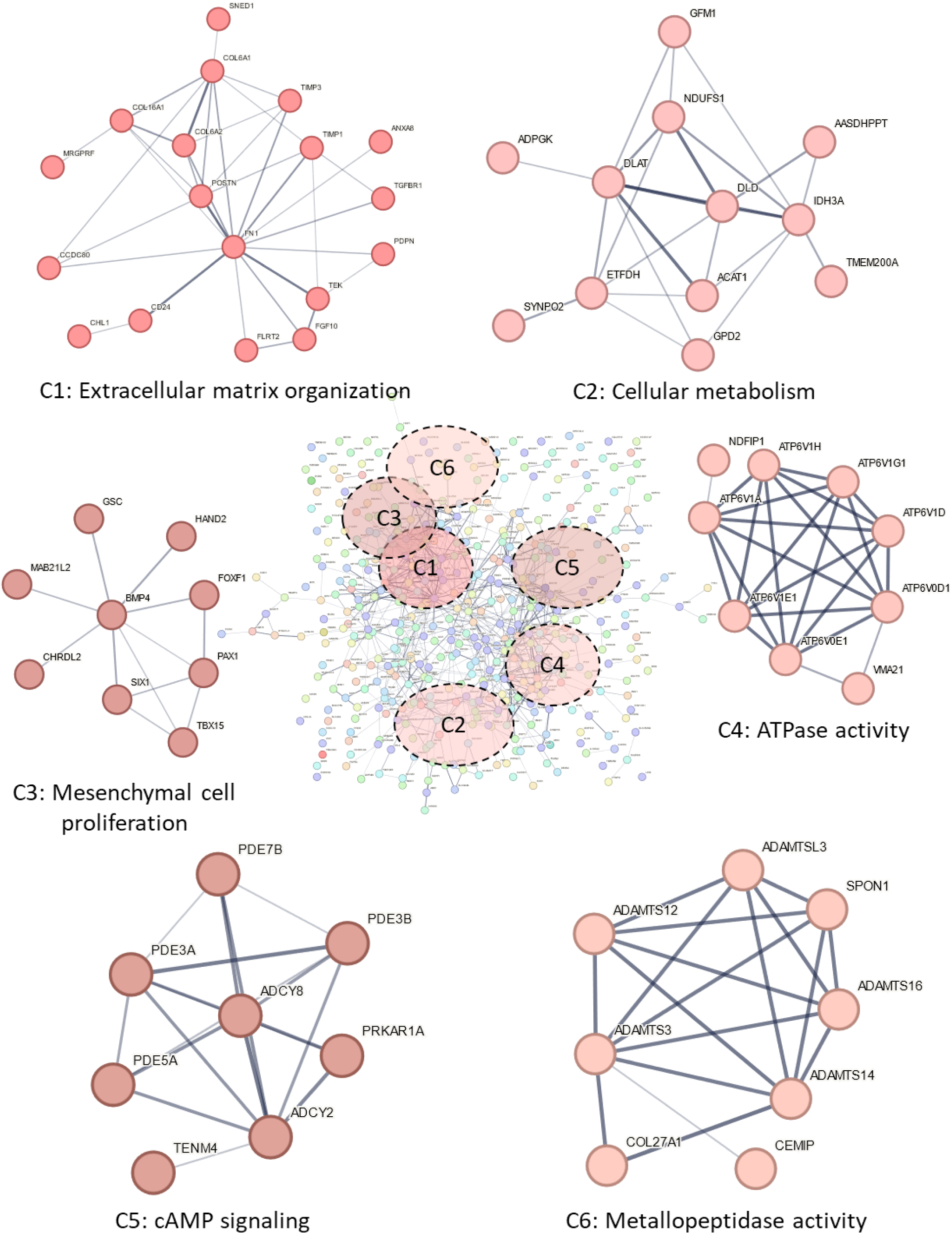
Pathways affecting several biological processes are enriched by genes found within the FD Signature. Using a statistical threshold of adjusted p<0.01, genes within the signature were loaded into the STRING database to determine potential protein-protein interactions of differentially regulated genes in FD. The most highly enriched processes involved “Extracellular matrix organization” (Cluster 1, C1), “Cellular metabolism” (C2), “Mesenchymal cell proliferation” (C3), “ATPase activity” (C4), “cAMP signaling” (C5), and “Metallopeptidase activity” (C6).

**Figure S2.**
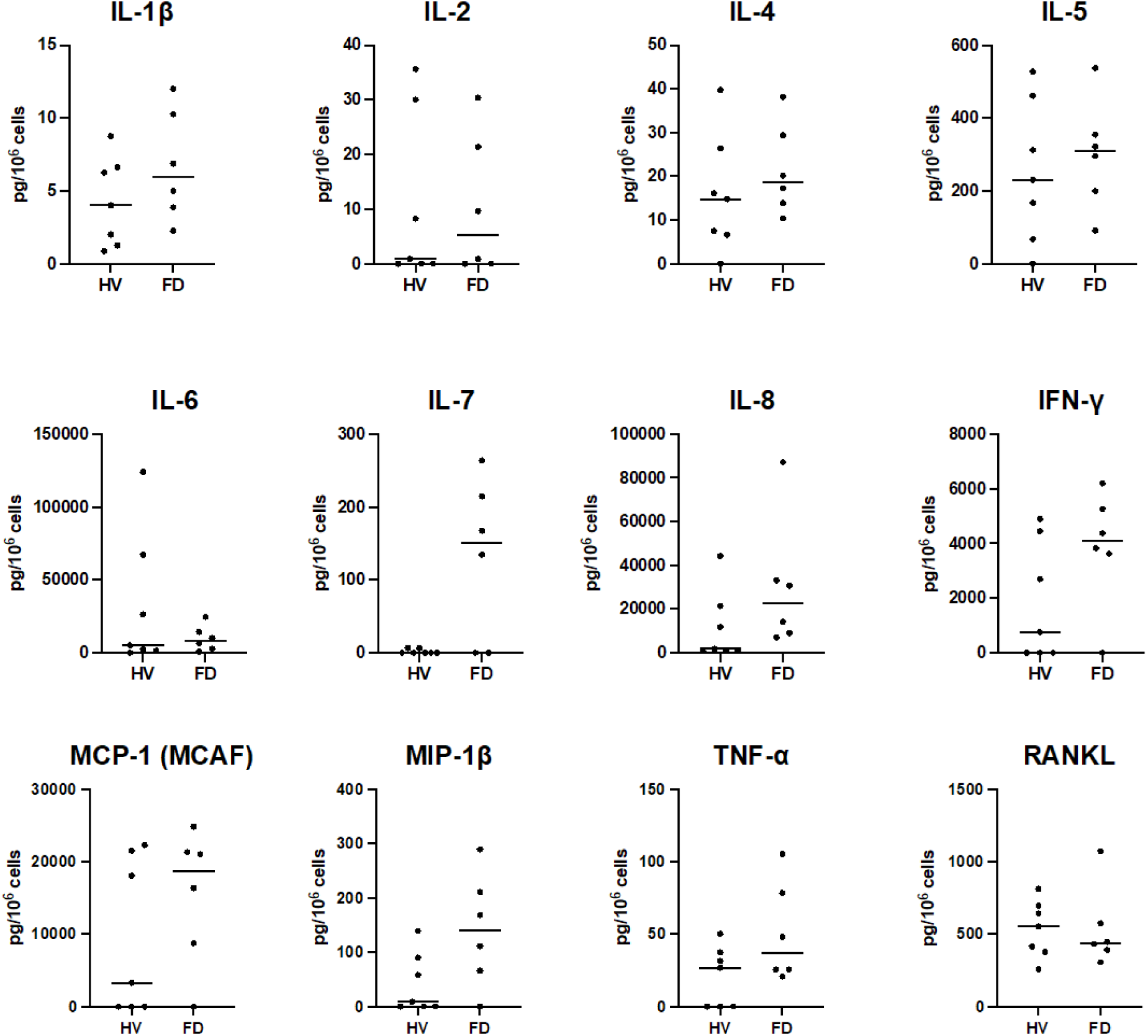
Cultured BMSCs derived from healthy volunteers (HVs) and patients with FD released pro-inflammatory cytokines, but differences were unable to be detected. Additionally, several factors were undetectable (IL-10, IL-12, IL-13, IL-17, G-CSF, and GM-CSF).

**Figure S3.**
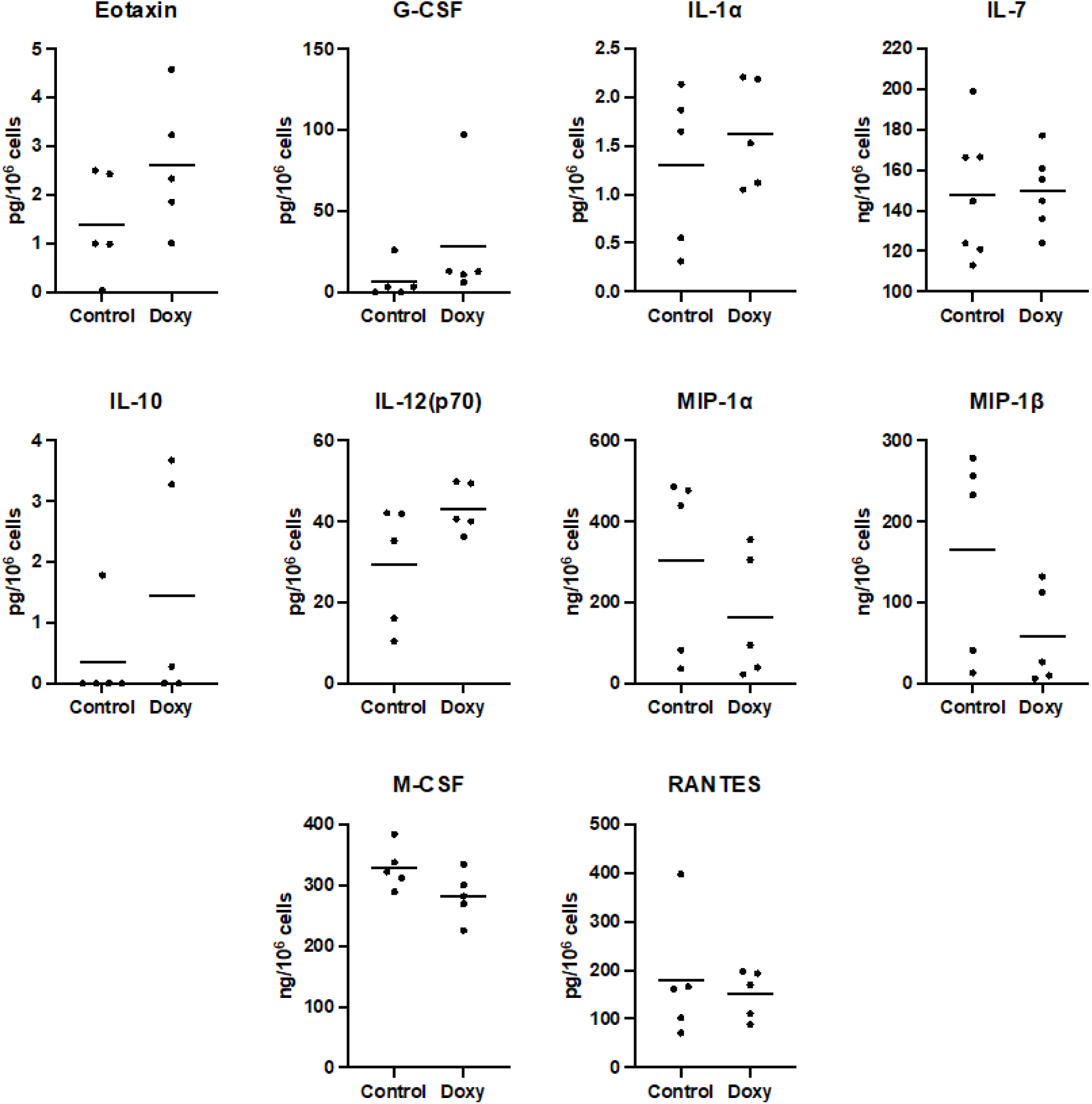
Additional cytokines expressed by cultured murine BMSCs. No differences were detected. IL-4, IL-9 were undetectable.

